# Trait-dependent species responses weaken the effects of response diversity on community stability

**DOI:** 10.64898/2026.06.26.734835

**Authors:** Anna Lena Heinrichs, Charlotte Kunze, Francesco Polazzo, Giulia Ghedini

## Abstract

The diversity of species responses to environmental change (response diversity) is a key mechanism of ecological stability. However, anticipating where strong or weak stabilizing responses emerge is challenging because species responses can depend on the local community and the specific stability metric. Whether species traits can consistently inform on how species respond to disturbances – enabling less context-dependent predictions – remains an open question. To address this gap, we use microcosm experiments on marine phytoplankton to test how response diversity supports multiple aspects of community stability under pulse temperature changes, testing both an increase (heatwave) and a decrease in temperature (coldspell). We then map species traits to their responses in a community to identify which traits modulate and predict species’ sensitivities. Fundamental response diversity, based on the diversity of species responses to temperature measured in isolation, was a weak predictor of community stability, and relationships differed between disturbances (i.e., heatwave and coldspell). Instead, species traits were consistent predictors of species responses in communities. Small, fast-growing species were more tolerant and benefited from the disturbance, while large, slow-growing species were less tolerant and decreased in proportion – these patterns were consistent across disturbances and community compositions. These results suggest that strong trait-performance relationships might reduce the importance of response diversity for stability. But these findings also show that general species traits, such as size and growth rate, can predict which species, and how, contribute to community responses, providing an empirical basis to relate species traits to stability outcomes under climate change.

## Introduction

Understanding the relationship between biodiversity and stability has been central to ecology for decades and remains a priority given ongoing climate change and biodiversity loss. More diverse communities are more likely to contain species that respond differently to environmental change (1–3): some species tolerate a disturbance or even benefit from it, while others suffer and decrease in abundance. The diversity of these responses, which has been termed *response diversity*, can generate asynchronous population dynamics that maintain a stable ecosystem functioning (4). Conceptually, it is well established that these stabilizing dynamics depend on the diversity and distribution of species traits within a community (5, 6) because, ultimately, traits influence the performance of organisms across environments (e.g., body size, physiology, life histories) (7, 8). However, the precise mechanisms through which biodiversity – either through variation in species responses or species traits – supports stability are still debated (9–12).

One issue is that trait-performance relationships are rarely linear and our understanding of how traits govern species interactions and performance in communities is incomplete (13, 14 (preprint 2026)). Given these difficulties, measuring changes in species performance over an environmental gradient can be a more powerful way to understand stability than just considering species traits. Changes in species performance can be used to quantify response diversity directly, which should be more closely related to community stability than traditional measures of diversity that do not incorporate performance (i.e., species richness or evenness) (15). However, empirical tests of how response diversity, measured on species performance, supports stability are few, and typically focus on a single aspect of stability (16–18). Since stability has many components (Box 1) (19–23), focusing on a single aspect limits a holistic understanding of how diversity buffers communities and might underestimate the risks of ecosystem change (20, 24–26).

Despite the appeal of measuring response diversity directly from species performance, understanding how species traits mediate the contribution of individual species to community stability remains essential for two reasons. First, obtaining performance-based metrics of response diversity is empirically demanding and does not offer a straightforward path if we seek predictions at global scales (27, 28). Second, traits may shape the degree of asynchronous or compensatory dynamics that stabilize communities, mediating the strength and effects of response diversity. In this context, Kunze et al. (29) proposed a novel framework to quantify how individual species contribute to changes in community properties under disturbance. This framework opens opportunities to identify the traits that make species more or less vulnerable to disturbances, which would increase our capacity to predict community stability under climate change.

A key feature of climate change is not only to increase mean temperatures but also the intensity and variability of temperature extremes (30, 31). Temperature is a powerful driver of ecological change because it affects growth, metabolism, and size – key traits linked to how species perform and interact (32, 33). In the microbial world, pulse changes in temperature can drive substantial community reorganization given the short life cycles of microbes (34, 35). Photosynthetic microbes, like phytoplankton, often reduce their cell size with warming (36–38). Given that cell size is linked to many aspects of organismal performance through its relationship with metabolic rate and nutrient uptake (39, 40), changes in phytoplankton cell size with temperature can affect the strength and stability of primary production at global scales (41).

We use phytoplankton as model system to H1) establish how response diversity drives five components of stability in response to pulse temperature changes (Box 1), and H2) test which traits predict species responses to these disturbances (Figure 1). We focus on five traits that directly reflect population performance (i.e., growth rate, thermal optimum, species dominance) or organismal functional traits that should underpin this performance (i.e., cell size and cell size plasticity) (H2, Figure 1). To test the generality of our predictions, we study these relationships under a pulse increase (Heatwave) or a decrease in temperature (Coldspell), representing a change in temperature of 5°C relative to control conditions (18°C) (see Appendix 1: Figure S1 for Disturbance set up).

### Box 1

**Definition of Stability metrics**

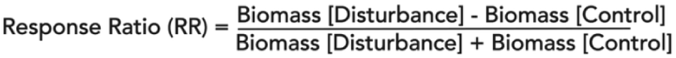

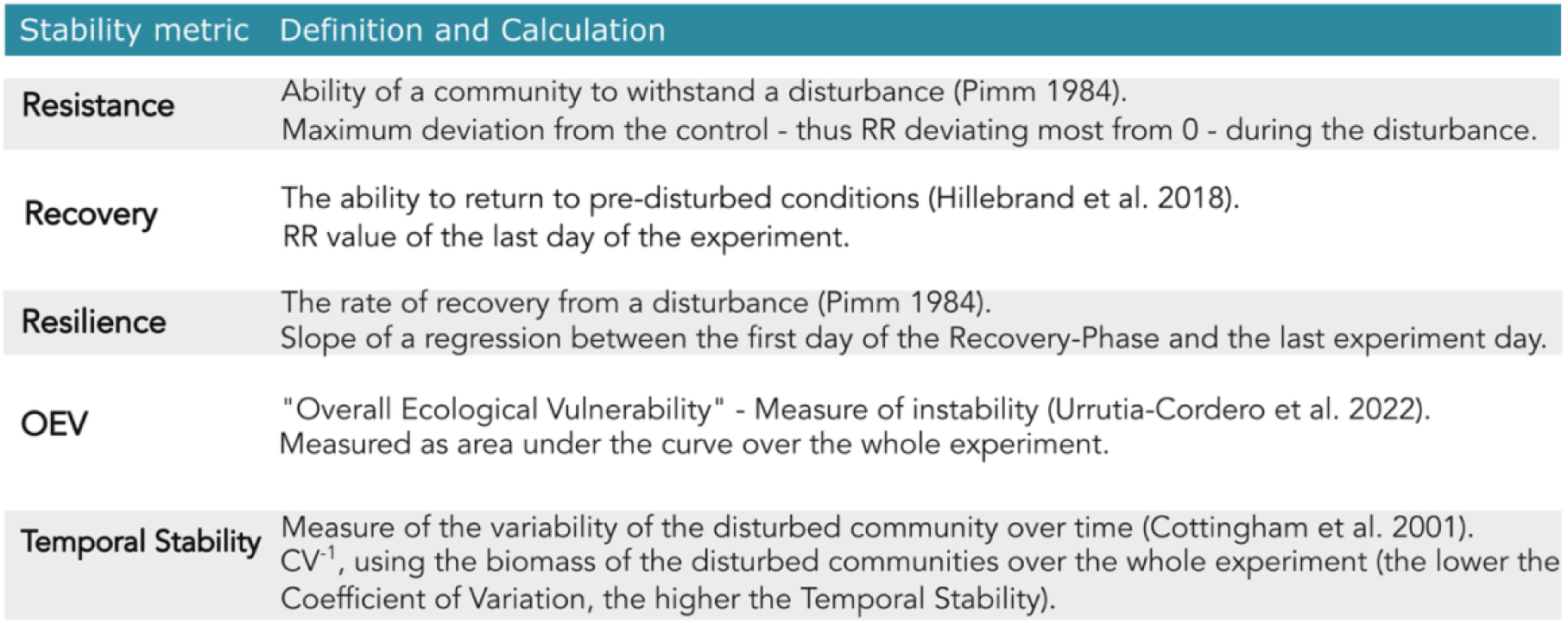

**Figure 1.**
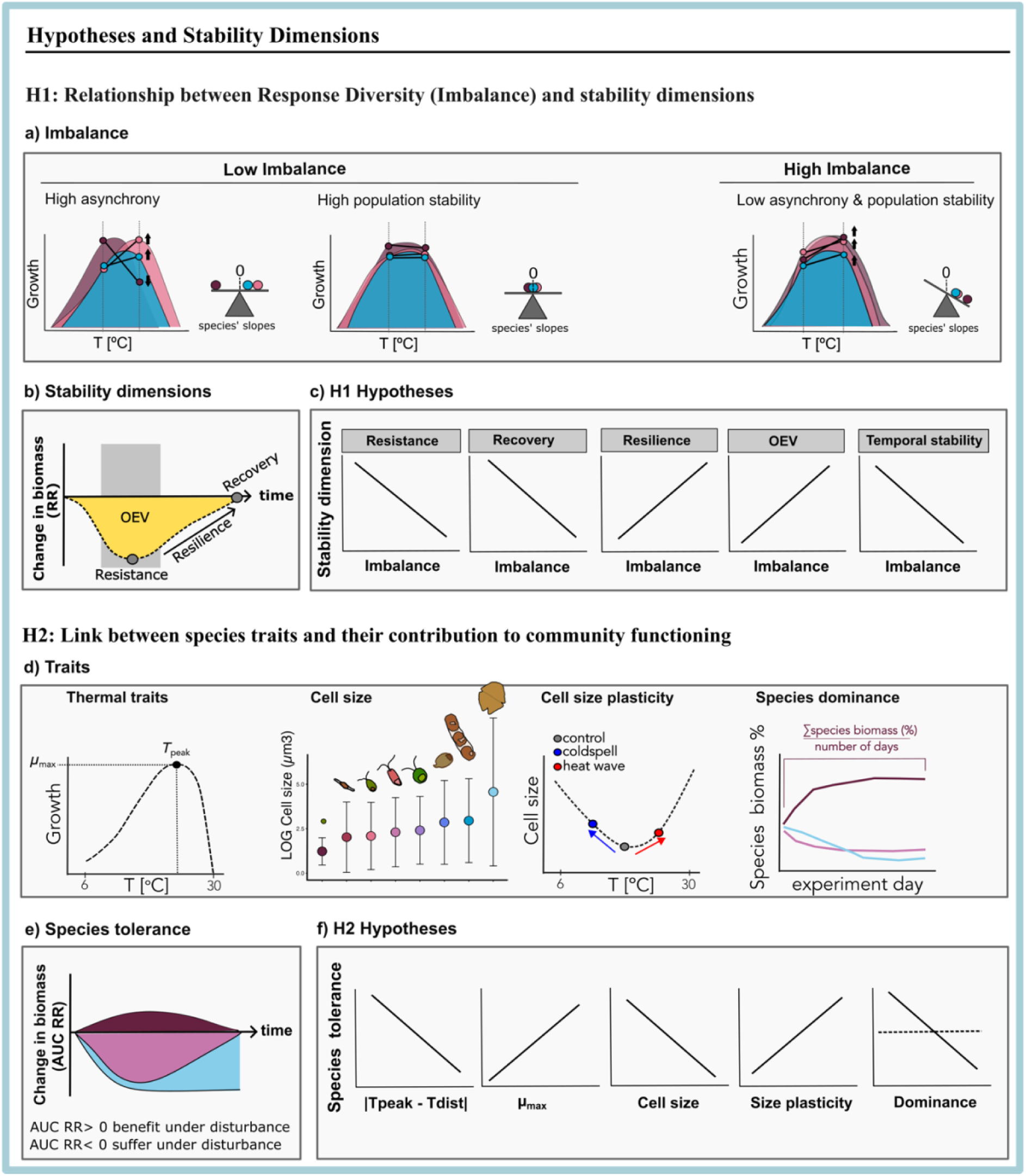
Conceptual figure of the Hypotheses (H1 & H2) and approach used to test each. To test the relationship between response diversity and stability under temperature change (H1), we manipulated response diversity using the imbalance metric (a). Imbalance is the mean of species responses (as an absolute measure) to a change in the environment (slope) measured in monoculture. A mean closer to 0 represents a more balanced community and values larger than 0 progressively more imbalanced communities. Stability metrics were calculated based on the Response Ratio (RR) of biomass between disturbed and control communities (b). c) Hypothesised relationship between each stability metric and imbalance. To test which and how species’ traits determine species’ tolerance/sensitivity to disturbances (H2), we determined species’ traits (d) by growing each species alone under a temperature gradient. Species tolerance (e) were calculated based on the area under the curve (AUC), using the RR of the species (the difference in relative biomass of each species between the disturbed and control communities) integrated over time. f) Hypothesised relationship between species traits and their sensitivity to disturbances.

### Manipulating response diversity using the imbalance metric

In each disturbance experiment (i.e., Heatwave and Coldspell), we manipulate the response diversity of phytoplankton communities, keeping the number of species constant. Our assessment of response diversity is based on the change in growth of phytoplankton species in monoculture along a temperature gradient (6-30°C), which was measured in independent assays prior to the disturbance experiments. Based on these assessments (thermal performance curves, TPCs), we estimate the change in performance (slope) of each species between the control temperature and the disturbance temperature (for each disturbance type; Heatwave and Coldspell). We then use the *imbalance* metric (17) to calculate and manipulate the response diversity of communities. Imbalance is calculated as the mean of species responses in a community, taking the absolute value (i.e., the mean slope). Based on this metric, we assembled nine communities differing in species composition but not richness (species richness = 3), creating a gradient of imbalance (response diversity) for each disturbance type. We then assess five stability metrics based on total community biomass, most of them calculated from the Response Ratio (RR) between disturbed and control communities (Box 1, Figure 1b).

Response diversity and imbalance are inversely linked: an imbalanced community has low response diversity because species respond similarly to the disturbance. Conversely, a balanced community typically has a high response diversity (where positive and negative responses offset each other), which should enhance stability; but a balanced community can also occur because species are insensitive to the disturbance (high population stability) (Polazzo et al. 2025). Therefore, the imbalance metric captures two mechanisms of stability linked to species traits: asynchrony in species responses to the environmental change, which should occur when response (and thus trait) diversity is high, and population stability caused by weak species responses, which might be tied to specific traits (Figure 1a).

This metric of response diversity, calculated from species responses in isolation, is termed *Fundamental Response Diversity (i.e., Fundamental Imbalance)*, and differs from the *Realized Response Diversity* which instead represents the actual variation of species responses in the community context. We focus on fundamental response diversity to test if community stability can be predicted solely based on changes in species’ performance measured in isolation, and thus independent of species interactions. We also test for the effects of realized response diversity to gain insights into the importance of species interactions in mediating the strength of response diversity. Fundamental response diversity has, to date, only been linked to community stability under fluctuating environmental conditions (17), but not under pulse disturbances, such as a Heatwave or Coldspell scenario.

## Results

### Response diversity is weakly related to community stability under pulse disturbances (H1)

We hypothesised that imbalanced communities, characterized by low asynchrony or low population stability, should be less stable under disturbances due to a lower chance of compensatory dynamics. We specifically predict that: imbalance should reduce community resistance and recovery, but increase resilience, given the often negative relationship between resistance and resilience (20, 42). We also expect that the overall vulnerability of communities should increase with imbalance and that their temporal stability should decrease (Figure 1b-c).

We found little support for these relationships. As predicted, higher imbalance (lower fundamental response diversity) reduced community *resistance* but only for communities exposed to the Coldspell (Figure 2a; slope = − 4.79, 95% CI [−7.9, −1.7]; Table S1). The trend was positive and not significant for communities exposed to the Heatwave. Imbalance was not correlated (or only weakly correlated) with the other stability metrics (Figure 2).

**Figure 2.**
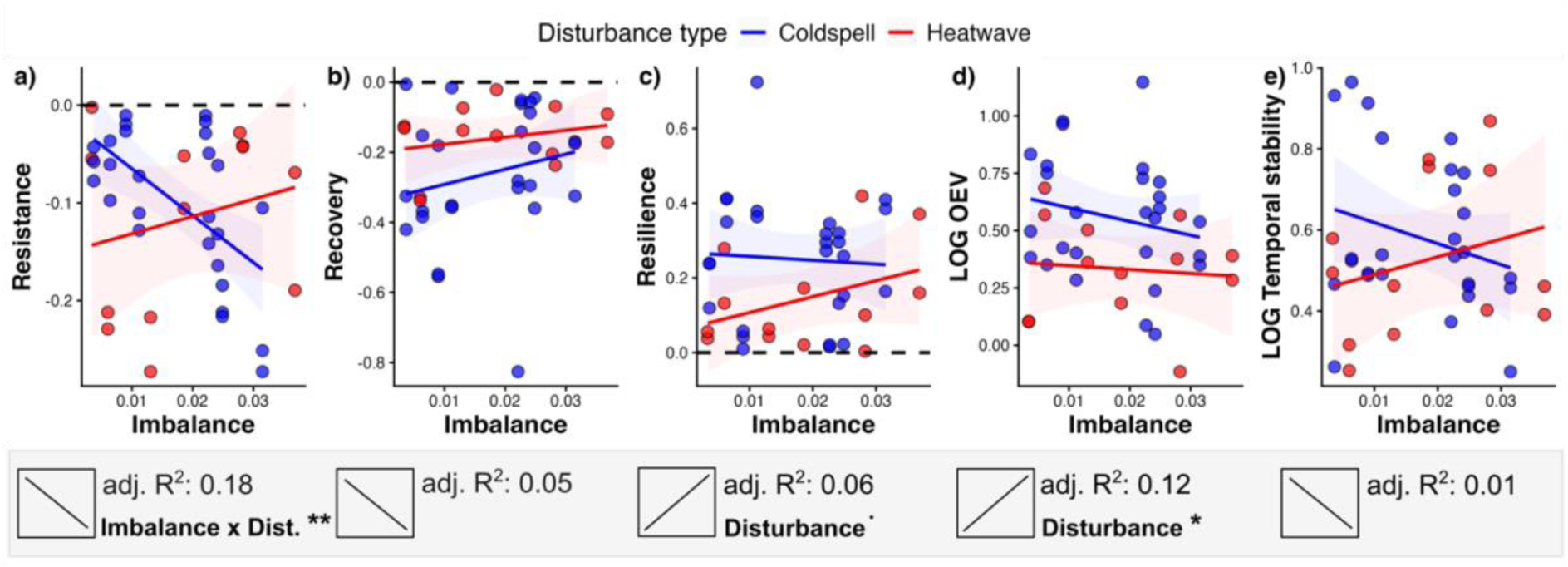
Relationship between stability dimensions, calculated on total community biomass, and imbalance under each disturbance type (*n* = 27 Coldspell in blue; *n* = 13 Heatwave in red). Imbalance correlated with only one stability dimension – resistance – and only under the Coldspell (a). There was no effect of imbalance on recovery, resilience, overall ecological vulnerability or temporal stability (b-e). Solid lines present the predictions from linear models, including imbalance and disturbance type as interactive predictors. Shaded areas show the 95% confidence interval. Boxes below each plot show the hypothesised relationships (H1, Figure 1), along with the adjusted *R^2^* from the model and significance values based on the ANOVA (^•^ p=0.05, * p < .01; ** p < .001; *** p < 0.0001). Temporal stability and OEV were log10-transformed to meet normality assumptions. Dashed horizontal lines in panels a) and b) indicate the maximum resistance and recovery (no difference between disturbed and control community, RR = 0), and in panel c) a Resilience of 0 (no change over time in RR).

The type of disturbance, i.e., the direction of temperature change, affected community stability in two cases. Heatwave-exposed communities showed lower *resilience* (F_1,36_ = 4.1, *p*-value = 0.05) and lower *Overall Ecological Vulnerability* (OEV; F_1,36_ = 6.4, *p*-value = 0.02) than Coldspell communities across the entire imbalance gradient, indicating a lower deviation from undisturbed conditions (Figure 2c; Table S1). We note that here resilience measures the rate of biomass change after the disturbance, without implying that communities return to pre-disturbed conditions.

Applying realized response diversity to our data, which represents the balance of actual species responses in the community, we found better support for the hypothesized relationship between resistance and imbalance: lower realized response diversity reduced resistance in both disturbance types. However, realized response diversity did not lead to more predictability of the other stability metrics (Appendix 1: Figure S2 and Equation S1).

### Species’ traits consistently underpin species’ tolerance to disturbances (H2)

To resolve which traits determine species responses to the disturbance, we integrated the difference in species proportional biomass change between the disturbed and control treatment over time (29). The area under the curve (AUC) of this change quantifies the species’ tolerance/sensitivity to the disturbance, where the species can either benefit (AUC.pi > 0) or suffer (AUC.pi < 0) (Figure 1e). Results based on absolute biomass change are the same and are shown in the supplements (see Appendix 1: Figure S3 for correlation between relative and absolute contribution).

We hypothesised that species are more tolerant to disturbances when the difference between their “optimal” temperature (temperature at which they grow fastest, *T*_peak_) and the disturbance temperature (*T*_dist_) is small (*T*_peak_-*T*_dist_ = 0). We also hypothesised that species with high maximum intrinsic growth rates (*µ*_max_) and small cell size will suffer less under the disturbance because these traits might allow faster recovery; and that greater size plasticity should also increase tolerance to the disturbance (because plasticity is linked to acclimation, independently of whether cells become smaller or larger). The role of species dominance might be disturbance- and trait-dependent, but generally, we expect a negative relationship between dominance and species tolerance, because rare species can benefit more from the decline of other, more abundant species (29) (Figure 1 H2).

Overall, the effects of traits on species’ tolerance were stronger and more consistent between disturbances than the effects of fundamental response diversity on stability.

As hypothesised, the ability of species to tolerate disturbances declined the further away their *T*_peak_ was from the disturbance temperature (Figure 3a; slope = − 0.07, 95% CI [−0.3, 0.2], ANOVA *T*_peak_: F_1,96_ = 16.14, *p*-value < 0.001; Appendix 1: Table S2). This was especially the case for the species exposed to the Heatwave, where the disturbance temperature consistently exceeded the temperature optimum of all species (see Appendix 1: Figure S4 for changes in growth between control and disturbance temperature; Table S3 for species’ *T*_peak_).

**Figure 3.**
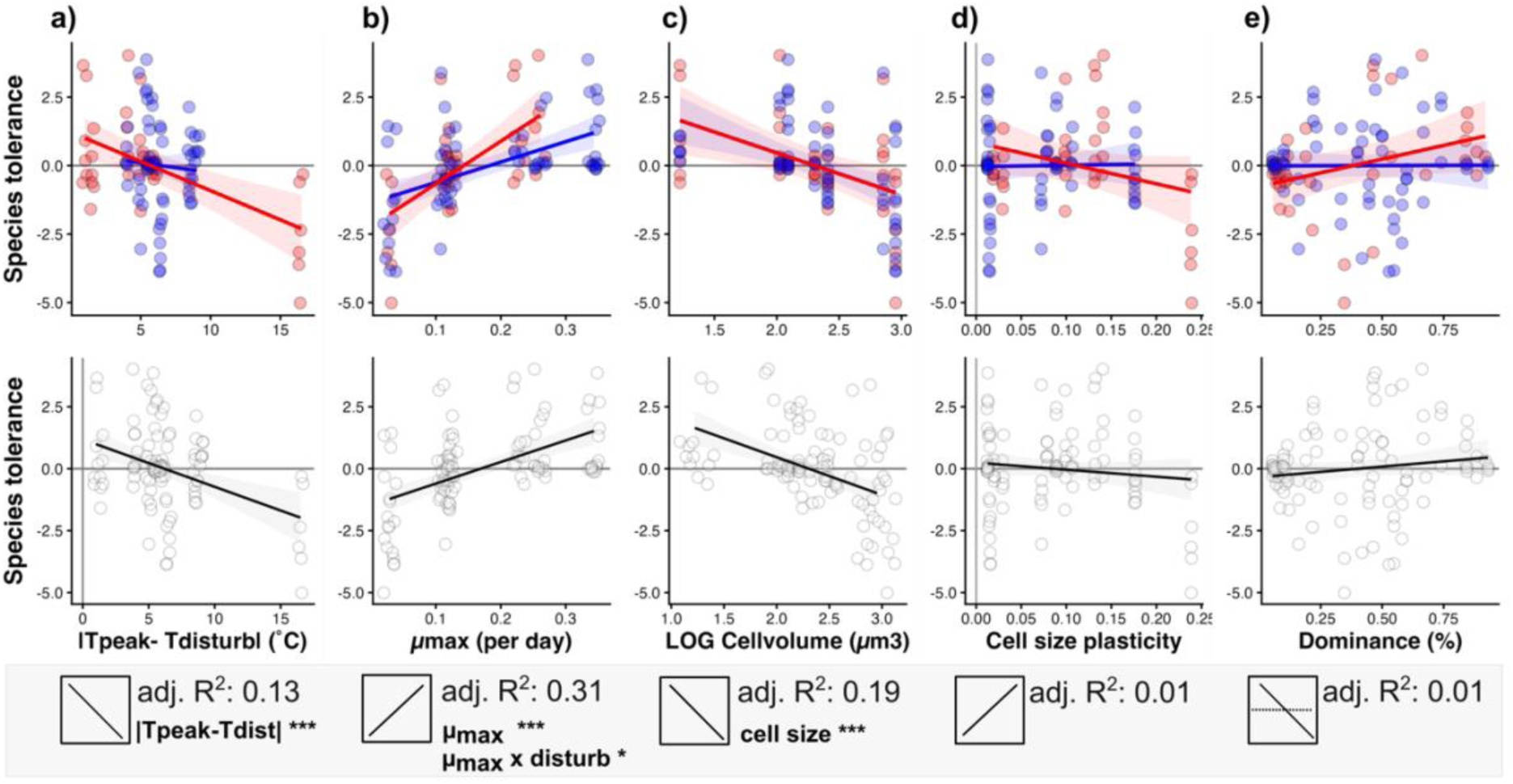
Linear relationships between species’ tolerance (in terms of relative biomass) and species’ traits (without the largest species *Levanderina*, see Results): (a) the absolute distance of a species’ *T*_peak_ and the disturbance temperature (*T_dist_*), (b) max. growth rates, (c) cell size under control conditions, (d) size plasticity as absolute cell size change from control to disturbance temperature, and (e) dominance. Upper row presents relationships testing for the effects of disturbance type (i.e., linear model including an interaction between the predictor and disturbance type) (Coldspell, *n* = 66; and Heatwave, *n* = 34). Lower row shows the general relationships between species traits and contribution, together for both disturbances (*n* = 100). Boxes at the bottom of each plot present the hypothesised relationships (H2, Figure 1), with adjusted R^2^ and significance values based on the ANOVA from the models in the top row (* p<.01; ** p<.001; *** p<.0001). Shaded areas show the 95% confidence interval.

Species with high maximum growth rates (µ_max_) suffered less than species with lower µ_max_ (Figure 3b). This positive relationship was stronger for communities exposed to the Heatwave (slope = 15.5, 95% CI [9.3, 21.8]) than for those exposed to the Coldspell (slope = 7.44, 95% CI [4.3, 10.5]), resulting in an interaction between *µ*_max_ and disturbance type (F_1,96_ = 5.3, *p*-value <0.001, Table S2).

Cell size (measured under control temperature conditions prior to the disturbance experiment) was negatively related to species tolerance, and its effect was consistent between disturbances (slope = − 1.5, 95% CI [−2.3, −0.7]; F_1,96_ = 26.8, *p*-value < 0.001). Smaller species benefited from the disturbance and increased in proportion compared to control conditions, while large species were most sensitive and decreased in proportion (Figure 3c).

Contrary to our expectation, cell size plasticity measured under the temperature gradient in isolation did not predict species’ tolerance to the pulse disturbances (Figure 3d).

Similarly, species relative dominance did not determine species sensitivity (Figure 3e). Even though there was a weak, but not significant, positive trend for the Heatwave communities: dominant species (under control conditions) had a higher tolerance to the Heatwave than rare species (slope = 2.0, 95% CI [−0.1, 3.9]).

### Community composition does not impact species’ sensitivity to disturbances

The effect of fundamental response diversity on stability varied between stability metrics and disturbance types. At a lower biological scale instead, traits more clearly determined how sensitive species were to disturbances in these communities. Here, we further test if species sensitivity were influenced by community composition and thus response diversity. We found that community composition played a minor role in determining how tolerant species were to disturbances (Figure 4) (ANOVA species’ composition, Heatwave: *F*_6,16_ = 1.9, *p* = 0.2; Coldspell: *F*_8,44_ = 3.8, *p* = 0.002). No species, except for *Nannochloropsis* (Na) in the Heatwave scenario, and *Chlamydomonas* (Chl) in the Coldspell, differed in their tolerance across community compositions (Figure 4).

**Figure 4.**
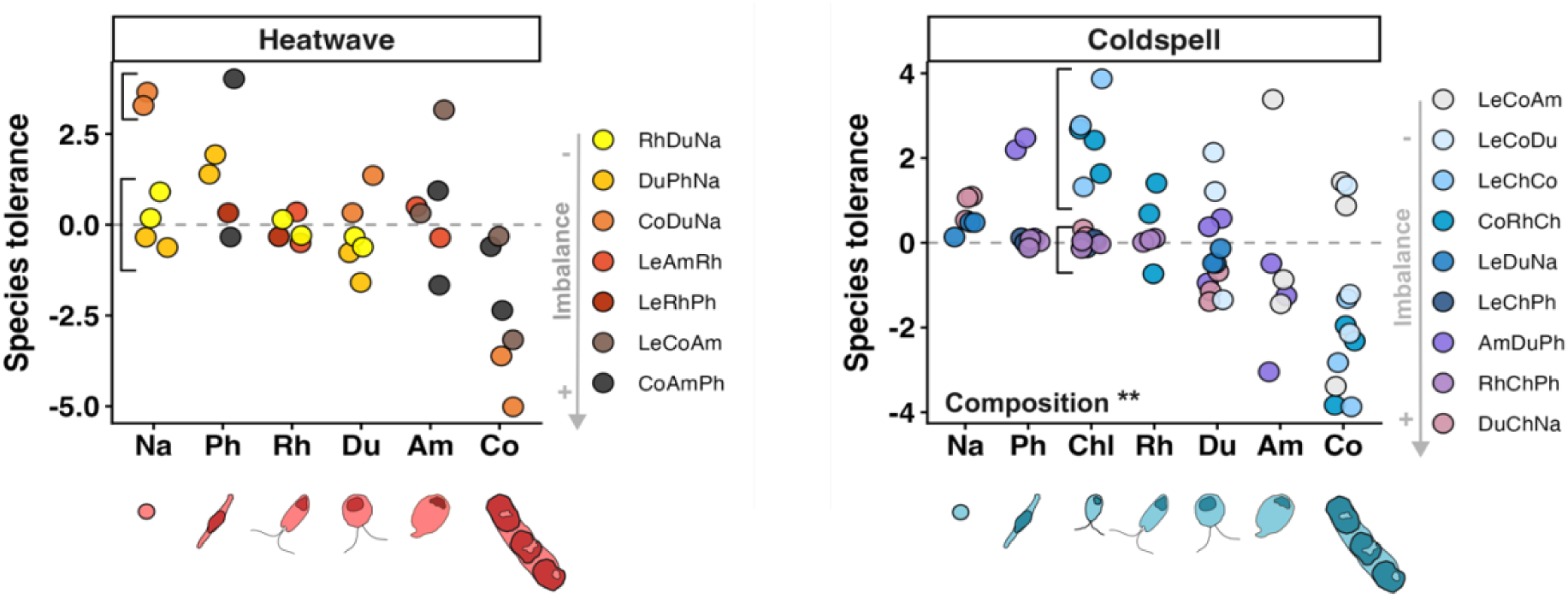
Species tolerance to disturbances across communities (differing in imbalance level and composition). For both disturbances, species tolerance mostly did not depend on the community they interacted with (the effect was non-significant for the Heatwave and significant for the Coldspell). But for each disturbance, composition affected the response of only one species, indicated by vertical lines (p < 0.05): *Chlamydomonas* (Chl) in the Coldspell and *Nannochloropsis* (Na) in the Heatwave. Species on the x-axis are ordered by cell size, from small to large (as shown in Figure 3c species tolerance declines with size). The large species *Levanderina* is excluded (see Figure S6 for data with *Levanderina*).

All results related to H2 exclude the largest species (*Levanderina*), which did not grow during the experiment (Appendix 1: Figure S5) and thus contributed little to community responses. We think that this species did not grow because it was introduced in communities at extremely low abundances, given its very large size (Table S3), experiencing stochastic dynamics unrelated to the disturbance. Results including this species show the same patterns (see Appendix 1: Figure S6 and Figure S7 for results with *Levanderina*).

## Discussion

Establishing a framework that connects species traits to community stability is a longstanding goal of ecology, and is a promising way forward to manage ecological systems under accelerating climate change and biodiversity loss (28). Conceptually, it is well established that species traits should influence response diversity and ultimately community stability. But the non-linearity of species responses to environmental change and their context-dependence create multiple hurdles in linking these ecological aspects. We empirically tested the predictive power of response diversity on community stability under pulse temperature changes, and then teased apart which traits drive these responses to mechanistically understand how species contribute to community change. Fundamental response diversity (based on the balance of species responses measured in isolation) was not a good predictor of community stability because it was affected both by the nature of the disturbance and the specific aspect of stability. Instead, general species traits consistently predicted species sensitivity to the disturbance and their contribution to community functioning, independently of the disturbance type (Heatwave and Coldspell) or community context.

Response diversity explicitly captures the diversity of species’ responses to the environment, in our case, changes in biomass growth with temperature (12, 17, 43). Therefore, we hypothesised that fundamental response diversity – the balance of positive and negative changes in species performance – would influence all aspects of community stability, with comparable effects between disturbances. To our surprise, resistance was the only stability metric (out of five) that was significantly related to fundamental response diversity (imbalance), and only under the Coldspell disturbance. Warm and cold conditions influence metabolism and growth in fundamentally different ways: cold temperatures generally slow metabolic processes, potentially reducing competition, whereas warming has the opposite effect which could reduce the strength of compensatory dynamics (44–47). Furthermore, our heatwave treatment was above the T_peak_ for most species, so the diversity of species responses might have been insufficient to stabilize communities under these extreme conditions. By measuring multiple aspects of community stability, our results suggest that fundamental response diversity only helps communities to resist, but not to recover, under pulse disturbances (the other stability metrics all depend on recovery (48)). Realized response diversity was a better predictor of resistance than fundamental response diversity – indicating that species interactions might change how species experience, and thus respond to a disturbance (18). But predictions did not improve for the other stability metrics that are a measure of recovery, suggesting that species interactions have smaller effects on how species respond after the disturbance. Therefore, it seems that response diversity does not determine the ability of a community to recover from pulse disturbances but presents an important property to predict community resistance.

The weak effects of response diversity on stability are unexpected – possibly because most theory is developed around fluctuating environmental changes (49) and because multiple dimensions of stability are rarely analysed simultaneously. Previous empirical work found that response diversity (imbalance) reduces the temporal variability of community functioning under environmental fluctuations (16, 17). Instead, imbalance did not stabilise macrophyte communities to pulse nitrate enrichment (50), similarly to our finding. Asynchronous species dynamics, which are at the core of response diversity, might work best under fluctuating conditions because they allow different “winners” and “losers”. Pulse disturbances instead are characterized by an abrupt change of short duration (Jentsch and White 2019), which shifts environmental conditions in a single direction before conditions recover (23). As the frequency and intensity of extreme events increase (30), understanding the mechanisms that stabilize communities under pulse disturbances becomes increasingly important (51).

Our analysis revealed that three traits best explained species’ responses to pulse disturbances: the thermal sensitivity of species measured as the distance between the optimal temperature and the disturbance temperature, their maximum growth rates measured over a temperature gradient, and average cell size measured under control conditions. All these traits were measured in monoculture and correlated with species responses in communities, so they can be considered “fundamental” traits rather than “realised” traits (similarly to the notion of fundamental *vs.* realised response diversity). Trait-driven species responses were independent of community composition: small, fast-growing species (*Nannochloropsis* and *Phaeodactylum*) always suffered less and increased in proportion than larger, slow-growing species under the disturbances (compared to control conditions). These results align with a recent study showing that the temporal stability of productivity in grassland species declines with increasing leaf area (52). Such (cell) size effect was not present in another study of phytoplankton communities under warming, where species’ relative contribution to community functioning was not related to species’ cell size (53). Smaller organisms generally have faster metabolic and growth rates than larger ones (37, 39, 54), which can facilitate rapid responses to changes in the environment and allow faster recovery, especially when the conditions after disturbance are favourable (55, 56). As such, organismal size is considered a good indicator of changes in community functioning under environmental change (40). Our results show that, despite the different effects of warm and cold conditions on metabolism, traits like cell size and intrinsic growth rates were consistent predictors of species responses between heatwave and coldspell disturbances.

Given the strong effects of T_peak_, µ_max_, and size on species sensitivity to disturbances, one might expect that these traits would influence species dominance in communities and, therefore, result in clear dominance-sensitivity relationships. However, there was no clear pattern in the sensitivity to disturbance of rare vs. dominant species, in line with previous work (29, 56–60). Overall, it seems that dominance rarely indicates the importance of a species for community stability. Dominance patterns might often be shaped by species interactions, whereas species traits might drive species responses to environmental change more directly.

Our work takes a step forward in mechanistically linking species traits to community stability. We show that response diversity only weakly correlates with stability because specific traits determine species’ sensitivity to the (pulse) disturbance. In other words, strong trait-response relationships at the species-level weaken the effects of response diversity on stability, explaining some of the inconsistent effects between studies. We identified three general traits that consistently predicted species responses to pulse temperature changes, independently of community composition and the direction of temperature change. Our findings suggest that small, fast-growing species will be key for stabilizing communities under rapid temperature changes. But this result is also worrying: pulse temperature disturbances can lead to a rapid re-organization of species hierarchies, which can make communities more susceptible to other types of environmental changes (61). Therefore, we highlight the importance of understanding the relative contribution of specific traits (e.g., growth rates, cell size) vs. trait diversity for stability outcomes – and the need to quantify how the importance of these processes varies with the type of environmental disturbance (press, pulse, fluctuating). Response diversity metrics, such as imbalance, capture mechanisms of stability that are driven both by the diversity of species responses and by population stability, which might be associated with specific traits, but do not straightforwardly allow to disentangle the two mechanisms. The combination of response diversity manipulations and trait-based analyses seems a promising way forward to better understand the mechanisms governing stability and anticipate which communities are more susceptible to climate change.

## Material and methods

### Overview

To test how response diversity (i.e., imbalance of a community) contributes to community stability, and how traits influence species sensitivity to disturbances (Figure 1), we conducted two separate experiments. First, we did a monoculture experiment (hereafter *Species’ responses experiment*) that provided the information on the responses of individual species along a temperature gradient (6°C − 30°C). These data were used to build Thermal Performance Curves (TPCs), that were used to calculate the imbalance of the communities (H1, Figure 1a), and to determine species’ thermal traits, cell size and its plasticity (H2, Figure 1d).

Then we ran a pulse disturbance experiment (hereafter *Disturbance experiment*) in which we exposed phytoplankton communities − consisting of different imbalance levels − to two disturbances, a Coldspell and a Heatwave. The communities were assembled with a fixed species richness (three species) using the species of the *Species’ responses* experiment, representing nine combinations of imbalance levels and community compositions. Below, we describe each experiment in detail.

### Species’ responses experiment

#### Experimental setup

We exposed eight marine phytoplankton species (monocultures), differing in size and taxonomic group (Figure 1d), to a temperature gradient to determine changes in growth rates and cell sizes with temperature. We used three replicates per temperature level (6**°**C, 10°C, 15°C, 18°C, 22°C, 25°C, 30°C), resulting in 21 units per species. For logistical reasons, we performed this experiment in four blocks of two species. The species were acquired from the Roscoff Culture Collection in France: *Nannochloropsis granulate* RCC8*, Phaeodactylum tricornutum* RCC2967*, Chlamydomonas sp.* RCC2607, *Dunaliella tertiolecta* RCC6, *Rhodomonas salina* RCC1506*, Amphidinium carterae* RCC88*, Coscinodiscus sp.* RCC4273*, Levanderina fissa* RCC10533.

Prior to the experiment, the species were grown at 18°C for at least 12 weeks in our laboratory. Species were then individually acclimated to the experimental temperatures for four days in dense cultures (∼ carrying capacity) before diluting them to equal cell abundance across acclimated populations (but different between species) that ensured a short and similar lag-phase within each species. The ideal cell abundance for each species was determined based on pilot experiments because temperature had large effects on species size, growth, and biovolume. Depending on species’ growth curves from the pilot experiment, start biovolumes of the different species ranged from 7,000 to 25,000 µm^3^/µl.

Before and during the experiment, the cultures were grown in F/2 media (62) in cell culture flasks with a volume of 160 ml (Sarstedt) at a light intensity of 70 µmol photons m^-2^ s^-1^ and a day:night cycle of 14:10. The same culturing conditions applied to the community experiment. The temperatures during the experiment were tracked using continuous data loggers (Hobo Pendant ®, Onset, Bourne, MA, USA).

To track species biomass and size over time, we took samples of 1 mL every other day and fixed them with Lugol’s Iodine Solution (4%). Cell abundance (cells µL^-1^) and cell size (major and minor axis in µm) were determined from pictures taken under a light microscope at 400x magnification (Olympus IX73 inverted microscope), and analysed using automated image analysis. We then calculated cell volume (µm³) based on the approximate geometric shape of each species (Hillebrand 1999) and determined the population biovolume as a proxy for biomass, as the product of cell abundance and cell volume. The experiment lasted until populations reached the stationary growth phase (between 21 and 30 days).

The intrinsic growth rates of each species were determined from changes in population biomass (biovolume) over time. Linear models were fitted to the exponential part of the growth curve on a logarithmic scale using the R package *‘growth rates’* and the *‘fit_easylinear’* function (Hall et al. 2014). For the samples exposed to extreme temperatures (i.e., 6°C and 30°C), the R package could not be applied for all species, as biomass declined over time. For those samples, we calculated the slope between the start and the end biomass using the following equation:

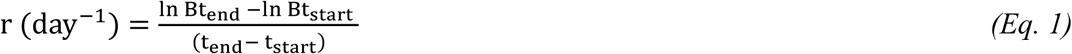

where *B* is the biomass at the end (t_end_) and at the beginning (t_start_) of the experiment.

#### Thermal performance curves

We used Generalized Additive Models (GAMs) for fitting Thermal Performance Curves (TPCs) based on species growth rates at each temperature. The TPCs allow us to calculate the community imbalance levels based on species variation in growth rates with temperature (see *Disturbance experiment* for details of the imbalance calculation) (Figure 1a) and to characterise two thermal traits for each species: *µ*_max_ as the maximum growth rate over the TPC range, and *T*_peak_ as the temperature at which *µ*_max_ occurs which are used to test H2 (Figure 1d, Appendix 2: Figure S8b).

#### Species traits

To link species traits to species’ sensitivity to disturbance (H2), we focus on two thermal traits (*T*_peak_ and *µ*_max_ as described above), and two morphological traits (cell size and cell size plasticity) (Figure 1d and Appendix 2: Table S3). We also calculated species dominance in control communities (this is explained in the community experiment).

The average cell size of species was estimated under culturing conditions (before the experiment started) at the control temperature (Figure 1d).

To determine plasticity in cell size, we fitted individual GAMs with cell size as the response variable and temperature as the predictor variable for each species, using cell size data from the exponential phase (Appendix 2: Figure S8a). We standardized cell sizes across all temperatures to make changes in cell size comparable across species:

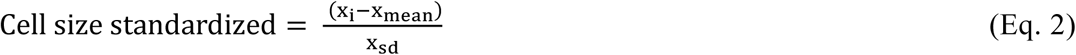

Where x is the cell size of each sample *i* within a species; x_mean_ is the mean cell size across all temperature treatments within a species, and x_sd_ the corresponding standard deviation.

We calculated plasticity as the slope between the standardized cell size at the control temperature and the cell size at the disturbance temperature used in the community experiment (i.e., 18°C to 23°C for the Heatwave, and 18°C to 13°C for the Coldspell) (Figure 1d). A positive slope indicates an increase in cell size from control to disturbance temperature, a negative slope indicates a reduction in cell size from control to disturbance temperature, and a value of zero means no plasticity (no change in cell size between control and disturbance temperature). We then took the absolute values to determine if greater size plasticity was predictive of species tolerance.

### Disturbance experiment

#### Experimental setup

To link response diversity to different stability metrics (H1) and test how species traits determine species’ sensitivity and their contribution to functioning under disturbance (H2), we conducted two 27-day pulse disturbance experiments. We used two disturbance regimes – a Heatwave and a Coldspell – to test the generality of our hypotheses. Each disturbance type represented an increase or decrease in temperature of 5℃ relative to control conditions, respectively. Since these two experiments were run separately, each experiment had its own control treatment.

For each experiment, we assembled 9 three-species communities that differed in response diversity (and thus composition) calculated as imbalance (explained below), using 3 replicates per temperature treatment (see Appendix 2: Table S4 for communities). Each of these 27 communities was either exposed to the control temperature or to the disturbance temperature (*n* = 54 communities per experiment). We had two incubators per temperature treatment and communities were randomly assigned between these two temperatures. All species in the communities started with the same initial biovolume of 30,000 µm^3^/µl, whereby all species had the same proportion in the community.

Each disturbance experiment (Appendix 1: Figure S1) began with a *Pre-Disturbance phase* of five days, during which all communities grew at the control temperature of 18°C. From day 6 to day 12 (*Disturbance phase*), the disturbed communities were either heated to 23°C (Heatwave) or cooled to 13°C (Coldspell), while the control communities remained at 18°C. It took five to eight hours until the disturbance temperature was reached. From day 12, the temperatures were regulated back to control (18°C) for two more weeks (*Recovery-Phase)*. Regulating temperatures back to control took around three to five hours (see Appendix 2: Figures S9 and S10 for temperature data). For the Heatwave experiment, one incubator assigned to the Heatwave disturbance heated up to 28°C during the night (instead of 23°C) for eight hours; these samples (14 samples of 27 samples in total) were removed from the analyses. Therefore, instead of nine imbalance levels, only 7 levels (and 7 communities differing in species composition) remained for the Heatwave experiment.

To track total and species biomass over time, we took 1 mL sample from each community every third day after homogenizing the samples carefully. The cell abundance, size, and biovolume of each species were calculated as described for the *Species’ responses* experiment. In each experiment phase, communities were refreshed with new media by removing 10 mL from each culture and replacing it with 10 mL of F/2 media. Because all species started with equal biovolume, the initial cell abundance of the largest species, *Levanderina fissas*, was necessarily low, as its cell volume was 40 times greater than that of the second-largest species. Throughout the experiment, its abundance remained consistently low (1–3 cell counts per sample), and no detectable population growth was observed (Appendix 1: Figure S5). To prevent strong fluctuations in community biomass arising from variation in counts of this large (Appendix 2: Table S3) and non-growing species, we calculated the mean biomass of *Levanderina* per sample for two experimental phases (Pre-phase & Disturbance phase, and Recovery phase).

#### Manipulating response diversity using the imbalance metric

To manipulate response diversity, we used species responses in monoculture to quantify the imbalance metric, which measures the variation in species’ responses in response to a change in the environment (17) (Figure 1a). Species responses are estimated as the change (slope) in intrinsic growth rate for each species between the control temperature and the disturbance temperature (see Appendix 1: Figure S4 for species’ slopes). Based on these slopes, we assembled species in different combinations to create 9 three-species communities that differed in Fundamental Imbalance (Figure 1a) – “fundamental” because it is based on species responses in monoculture – calculated as:

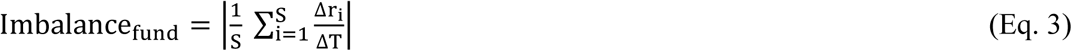

Where S is the total number of species in a community (3 in our case), Δr the change in intrinsic growth rate of species *i*, and ΔT the temperature change (between control and disturbance temperature, always 5°C in this study). Thus, fundamental imbalance is the mean of the slopes between all species in a community, such that values of 0 indicate a balanced community, either because of high variation in species responses (asynchrony – directly related to response diversity) or high population stability (all species respond minimally to the environmental change) (Figure 1a).

The resulting fundamental imbalance gradient ranged from 0.003 to 0.038 for the Heatwave experiment and from 0.004 to 0.032 for the Coldspell (Appendix 2: Table S4). The imbalance metric does not have finite range (as it depends on the species and variable measured; see Appendix 1: Figure S2 and Equation S1 for calculation of realized Imbalance).

#### Stability metrics

We tested how response diversity affects multiple stability dimensions: resistance, recovery, resilience, Overall Ecological Vulnerability (OEV), and temporal stability (Figure 1a and Box 1). All stability metrics, except temporal stability, were determined based on the difference in community biomass between disturbed and control communities using the following equation:

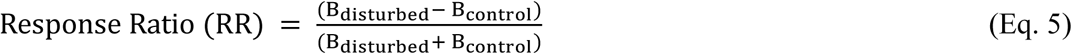

where *B_disturbed_* is the community biomass of the disturbed communities, and *B_control_* the community biomass of the control communities at a given time point (i.e., each sampling day) (see Appendix 2: Figure S11-S12 for community biomass data during the experiments). RR varies between −1 and 1 for each time step, where a positive RR indicates higher community biomass in the disturbed communities, while a negative RR indicates higher community biomass in the control communities.

*Resistance*, which describes a community’s ability to withstand a disturbance (19), was calculated based on the maximum absolute deviation in community biomass from the control during the disturbance phase (day 6, 8, and 11) (63).

*Recovery*, which describes the ability to return to pre-disturbed conditions, was defined as the RR at the last day of the experiment (day 27) (20). For both resistance and recovery, we defined all deviations from the control as destabilizing, regardless of their sign (i.e., whether they were driven by an increase (positive sign) or decrease (negative sign) in biomass), so we converted all positive deviations (positive RR values) to negatives (Figure 1b); thus, a value of 0 means maximum resistance or full recovery.

*Resilience,* which describes the rate of recovery from the disturbance (19), was calculated as the slope of a regression between the RR of the first day after the disturbance (day 12) and the last day of the experiment (during the recovery phase) (Figure 1b). We note that most of our communities did not fully recover after 27 days (RR≠ 0), therefore a higher resilience does not mean that the community returns to pre-disturbance conditions, just that its biomass changes after the disturbance at a faster rate. Similarly to above, we converted all slopes to positive values to have higher rates of change associated with steeper slopes – independently of whether the disturbance caused an increase or decrease in biomass.

The *Overall Ecological Vulnerability* (OEV) is a measure of instability and was calculated as the absolute area under the curve of the difference between disturbed and control community biomass over the whole experiment (21) (Figure 1b). The larger the OEV (absolute area), the more the disturbed communities deviated from control, indicating a higher vulnerability. *Temporal stability*, which is a measure of how stable a community process is over time, was calculated as the inverse of the Coefficient of Variation (CV), based on the total biomass from all days of the disturbed communities. A higher CV indicates greater temporal variability in community biomass (1, 19, 63); therefore, temporal stability increases with lower CV values (64).

The Response Ratio of the pre-phase was used as baseline and subtracted from the Response Ratio determined over the whole experiment, to exclude that differences between control and disturbed communities not caused by the disturbance itself influence the stability metrics.

#### Species tolerance to disturbances

To test whether species’ traits determine species sensitivity to environmental change (H2), we build on Kunze et al.’s (2024) approach, which transfers the integrative metric of the area under the curve, previously used for communities (OEV) (21), to the species level (AUC_sp_) (29). Thereby, the difference in species biomass between the disturbed and control treatment is integrated over time, using Equation 5, but using either species absolute biomass (RR) or relative biomass (Δpi) instead of community biomass (Figure 1e). This approach shows species’ tolerance to environmental change, based on species absolute changes in biomass (absolute tolerance) and relative to other species based on species proportions (relative tolerance). The sign of the species’ AUC (positive or negative) and the absolute or relative tolerance define whether the species benefits from the disturbance by increasing in biomass and proportion, or suffers from it by declining in both metrics, or the alternative combinations (see Kunze et al. 2024 for details).

As for the community Response ratio, we used the pre-phase RR again as baseline and subtracted it from the species RR determined over the whole experiment.

#### Relative dominance

In addition to the four traits derived from the *Species’ responses* experiment, we examined whether species’ tolerance to disturbances is associated with their dominance in the community. To determine species dominance, we averaged species proportional biomass (%) within the control community during the experiment, whereby high and low proportions present dominant and rare species, respectively.

#### Statistical analyses

To test the relationship between stability dimensions and imbalance (H1), we used linear models with imbalance and the disturbance type (Coldspell an d Heatwave) as interactive predictors (*n* = 40). From the linear model, we extracted the slope estimates as well as R^2^ and used ANOVAs to test for the significance of the relationships. To ensure normality of residuals, we log-transformed OEV and temporal stability. For temporal stability, we excluded one outlier in the Heatwave communities (*n* = 39). Note that 14 of 27 samples had to be removed from the Heatwave experiment due to a defective incubator, resulting in 13 instead of 27 samples in total for the Heatwave experiment.

To test the relationships between species relative tolerance and species traits (H2), we used again a linear model with the traits and disturbance types as predictors, including an interaction term (*n* = 100). From those linear models, we extracted the slope estimates and R^2^ and tested for the significance of these relationships via ANOVA. We used the relative AUC (AUC p_i_) that excluded the large species *Levanderina* that, although it did not grow during the experiment, fluctuated randomly over time which influenced species proportions strongly, given its large cell size (see Appendix 1: Figure S6 and S7 for data with *Levanderina* with *n* = 120). Lastly, to test how species’ tolerance change across community compositions, we used an ANOVA for each disturbance experiment with species ID and composition as interactive effect (*n*=34 [Heatwave]; *n*=66 [Coldspell]), and significant differences across communities were checked via Tukey Honest Significant test.

## Acknowledgements and Funding

We thank Ricardo Estevens for supporting sampling during the experiment and Samuel R. P.-J. Ross for his helpful comments on an earlier draft of the manuscript. The work was supported by an ERC Starting Grant to GG (project META_FUN, 101116029). Views and opinions expressed are however those of the author(s) only and do not necessarily reflect those of the European Union or the European Research Council. Neither the European Union nor the granting authority can be held responsible for them.

## Author Contributions

**Anna Lena Heinrichs:** Conceptualization (equal); Data curation (lead); Formal analysis (lead); Investigation (lead); Methodology (equal); Validation (equal); Visualization (lead); Writing – original draft (lead); Writing – review and editing (equal); **Francesco Polazzo:** Conceptualization (equal); Formal analysis (supporting); Methodology (equal); Validation (equal); Writing – review and editing (equal); **Charlotte Kunze:** Conceptualization (equal); Formal analysis (supporting); Methodology (equal); Validation (equal); Writing – review and editing (equal); **Giulia Ghedini:** Conceptualization (equal); Data curation (supporting); Formal analysis (supporting); Funding acquisition (lead); Methodology (equal); Project administration (lead); Supervision (lead); Validation (equal); Visualization (supporting); Writing – original draft (supporting); Writing– review and editing (equal).

