## Appendix for "Trait-dependent species responses weaken the effects of response diversity on community stability"

**Corresponding authors:** Anna Lena Heinrichs and Giulia Ghedini

**Appendix 1:**

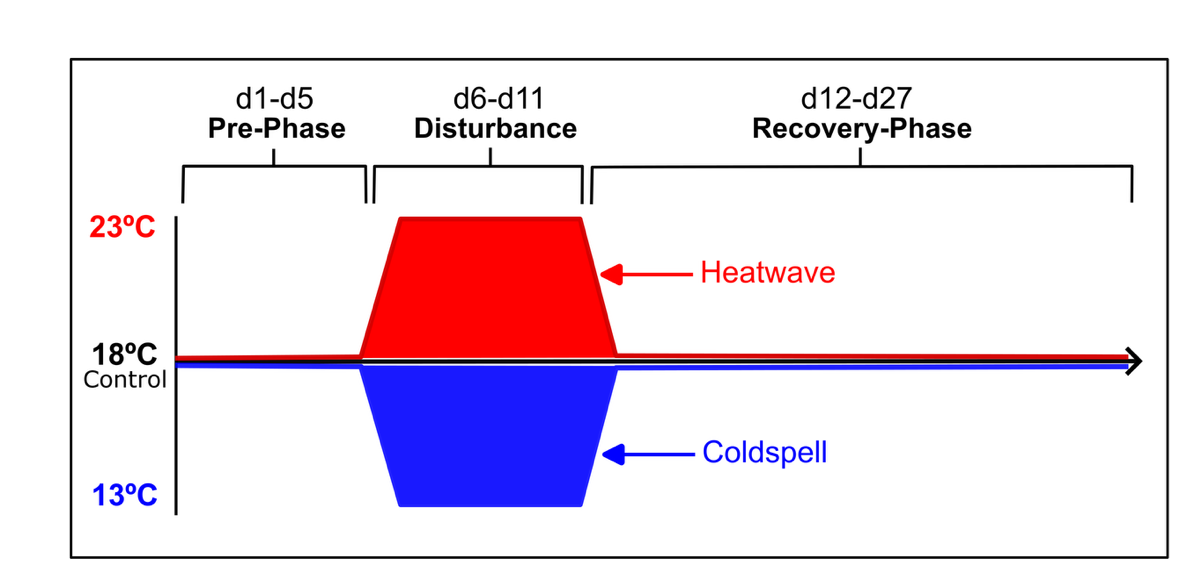

**Figure S1.** Experimental setup of the Disturbance experiment. Control temperature was 18˚C. For the Heatwave, the temperature was heated up to 23˚C, and for the Coldspell, the temperature was regulated down to 13˚C. The experiment started with a 5-day Pre-Phase where all communities were kept at the control temperature. The disturbance phase was from day 6 to day 11, where the disturbed communities were either heated up (Heatwave) or cooled (Coldspell). From day 12 onwards, there was a Recovery-Phase during which the disturbed communities were returned to temperature control until the end of the experiment (day 27).

**Realized Response Diversity** was calculated as the mean response of species in a community to the disturbance, using the mean of the AUC RR of all species in the community. Thereby, the difference in species biomass between the disturbed and control treatment is integrated over time, using Equation 5, but using species absolute biomass (AUC RR):

$Realizid Response Diversity= \frac{1}{S} \sum_{i=1}^{S} AUC(\frac{\mathrm{biomass}\left[ \mathrm{disturbed} \right]-biomass \left[ \mathrm{control} \right]}{\mathrm{biomass}\left[ \mathrm{disturbed} \right]+biomass \left[ \mathrm{control} \right]})$	(Eq. S1)

where: *S* presents the total number of species in the community.

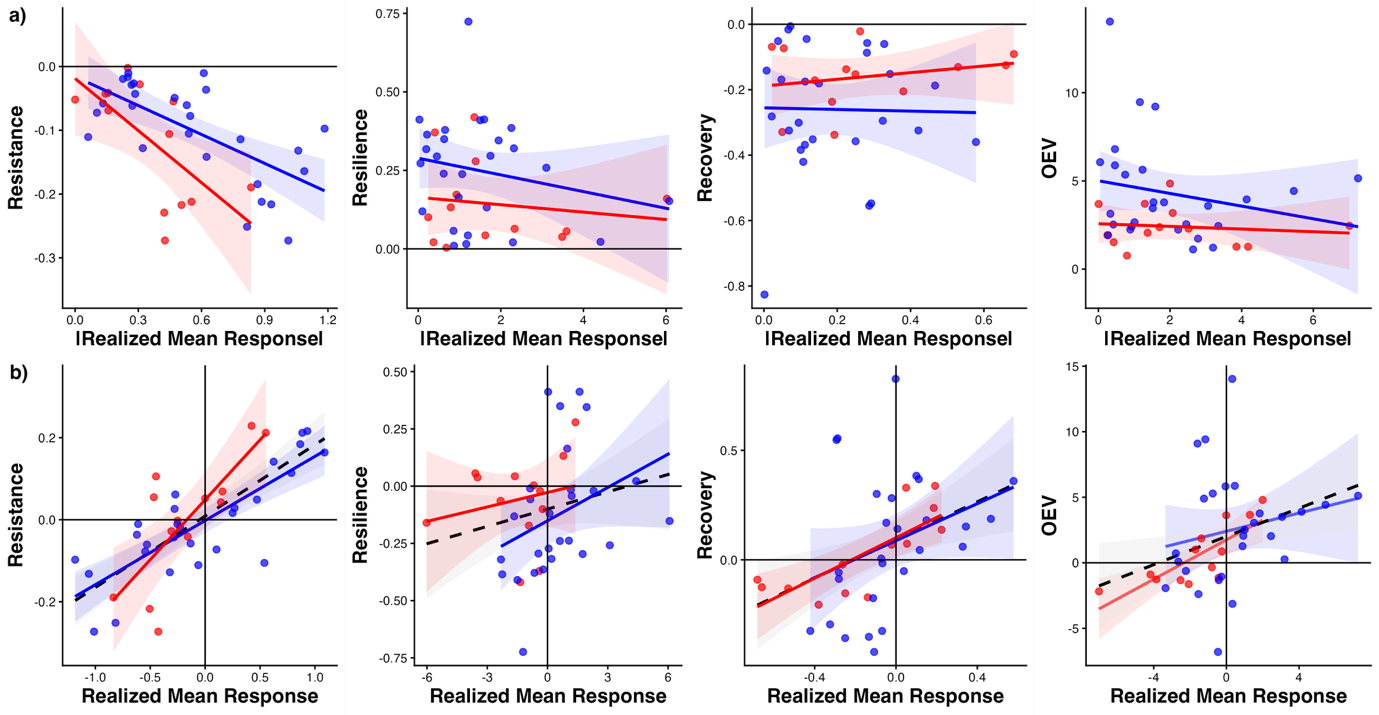

**Figure S2.** Realized Response Diversity measured as the Mean Realized Response (Eq. S1). A realized mean response of 0 means a maximum balanced community. **a)** Effects of absolute realized Imbalance on stability metrics across disturbance types (blue=Coldspell; red=Heatwave). Only resistance showed a dependency, while other metrics showed no relationship to Realized Mean Response. **b)** When considering the direction of destabilization, we recover the expected positive relationship between community biomass change and the mean biomass change of species in these communities, according to Kunze et al. (2026) (positive=higher biomass than control community; negative=lower biomass than control communities, zero=maximum stability as control equals disturbed communities).

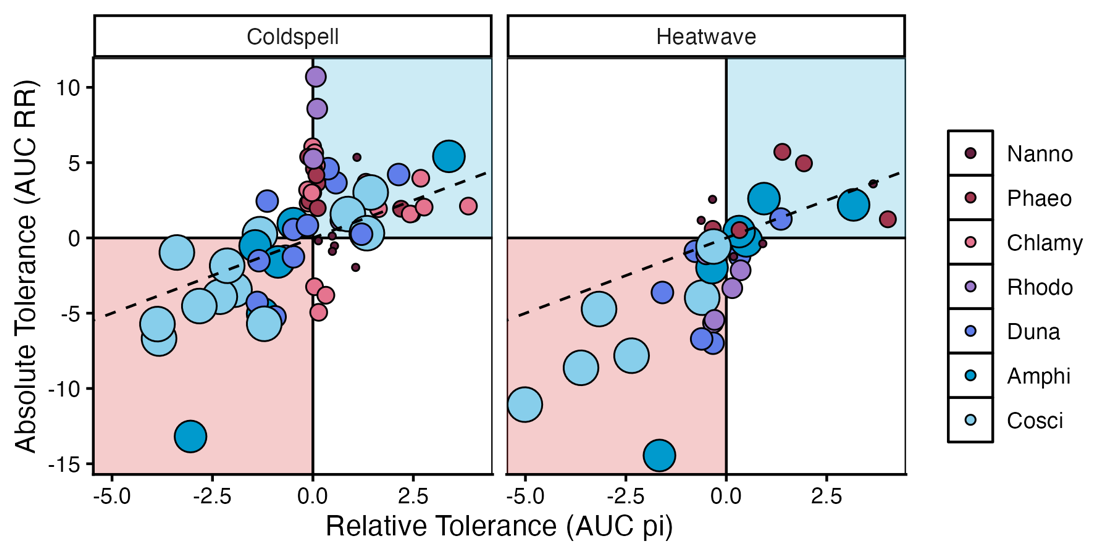

**Figure S3.** Relative species tolerance vs. Absolute tolerance, without the large species Levanderina. The size of data points reflects species’ cell size.

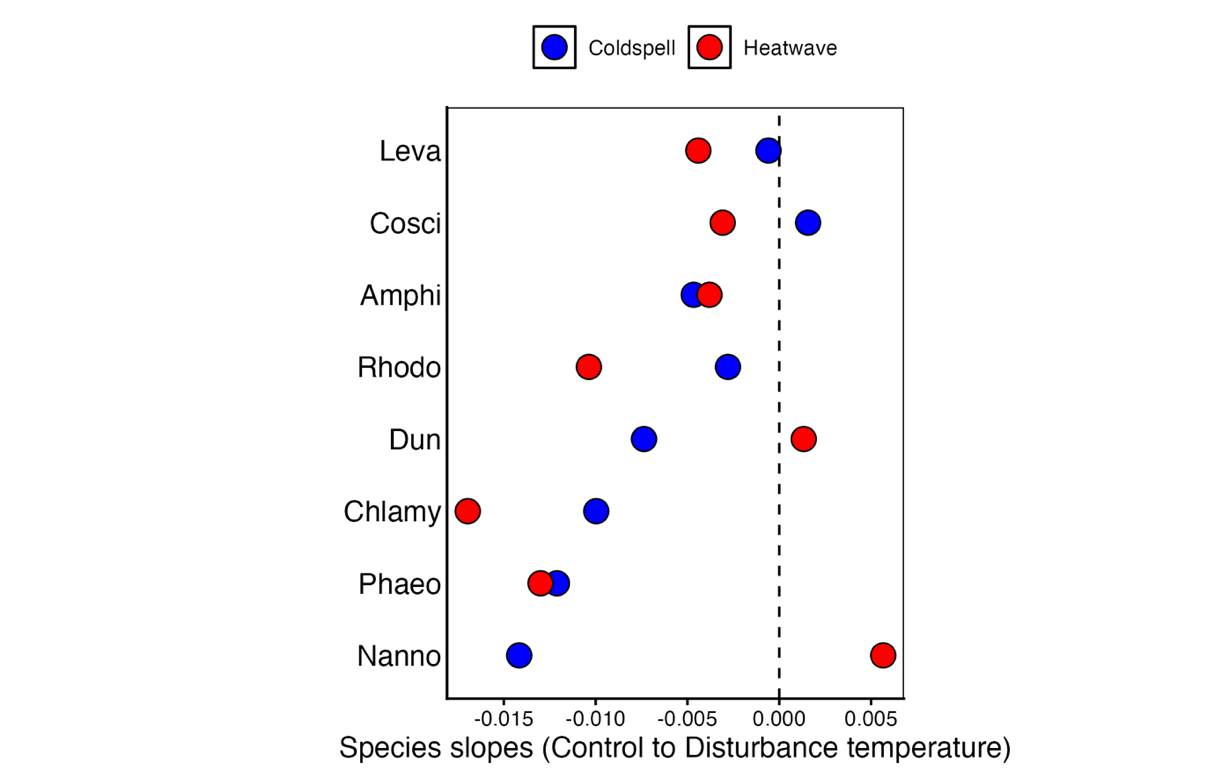

**Figure S4.** Species slopes based on growth rates between control temperature and disturbances temperature, accordingly to the TPCs in Figure S8. Slopes were used to calculate the Imbalance levels of communities. Colours present the disturbance temperatures of each experiment (blue = Coldspell, red = Heatwave). Species are ordered by cell size from small to large, with Nanno (bottom) being the smallest.

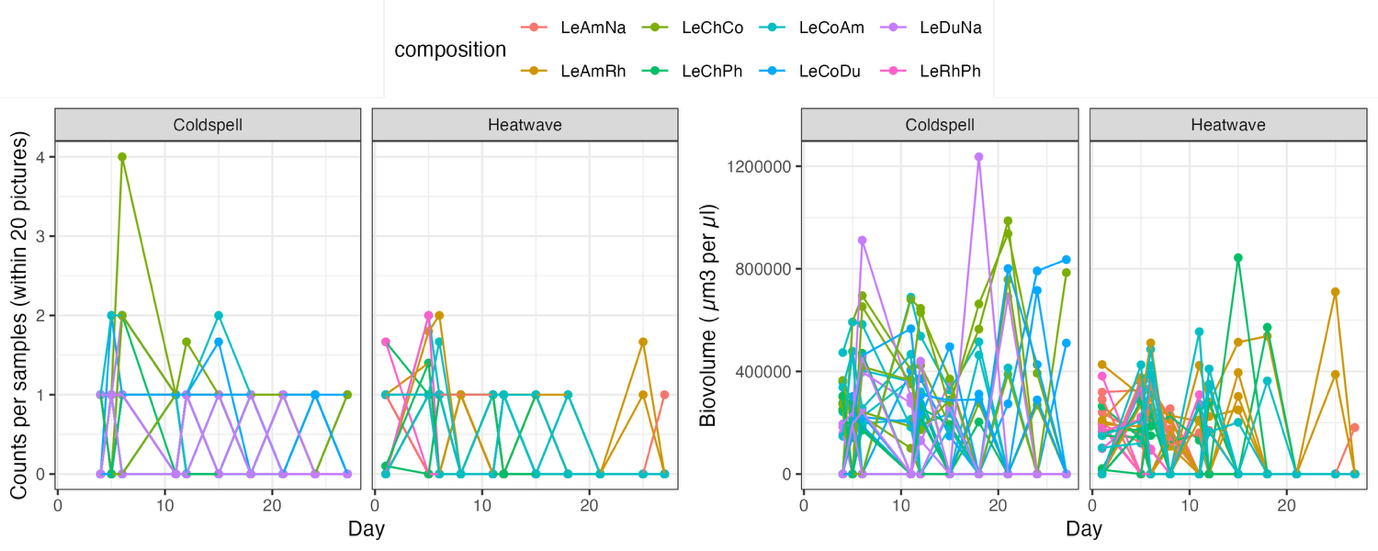

**Figure S5.** Counts and biovolume data of the largest species Levanderina during the experiments (i.e., Coldspell and Heatwave) show that there was daily variation in abundance but no clear trend over time.

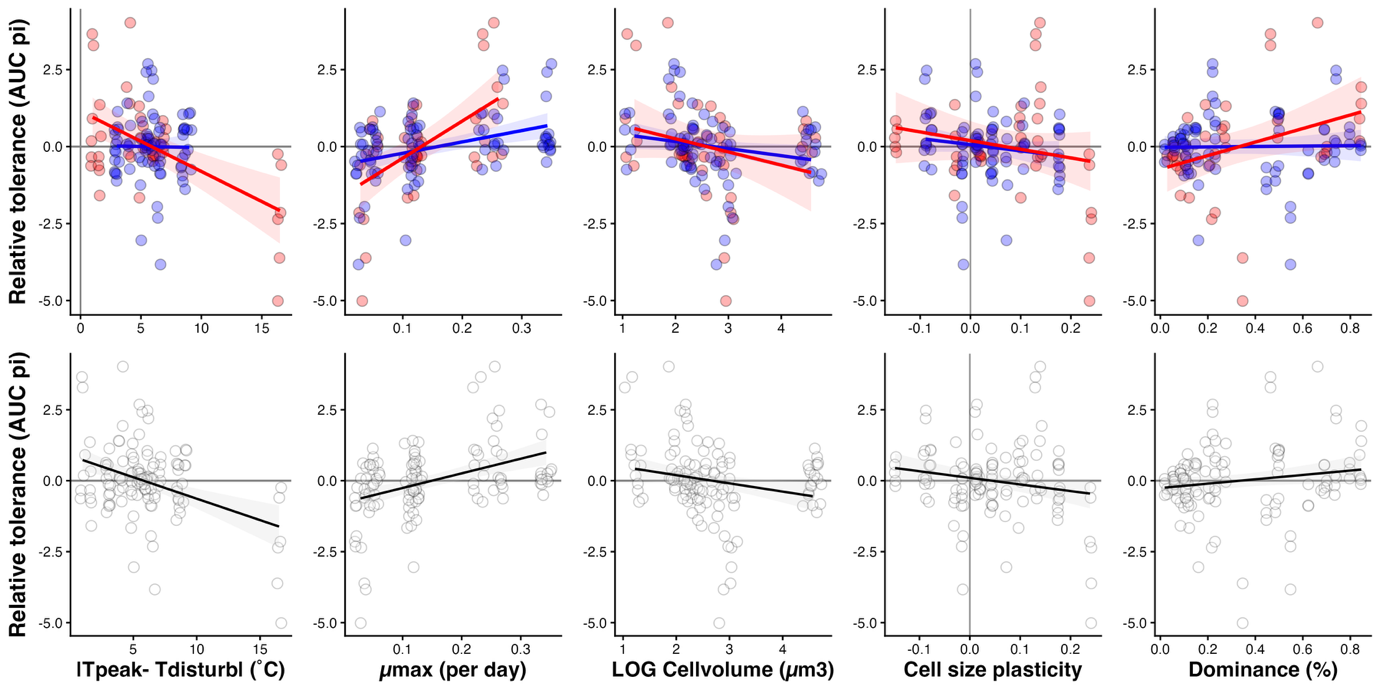

**Figure S6.** Species relative tolerance vs species traits, including the large species Levanderina.

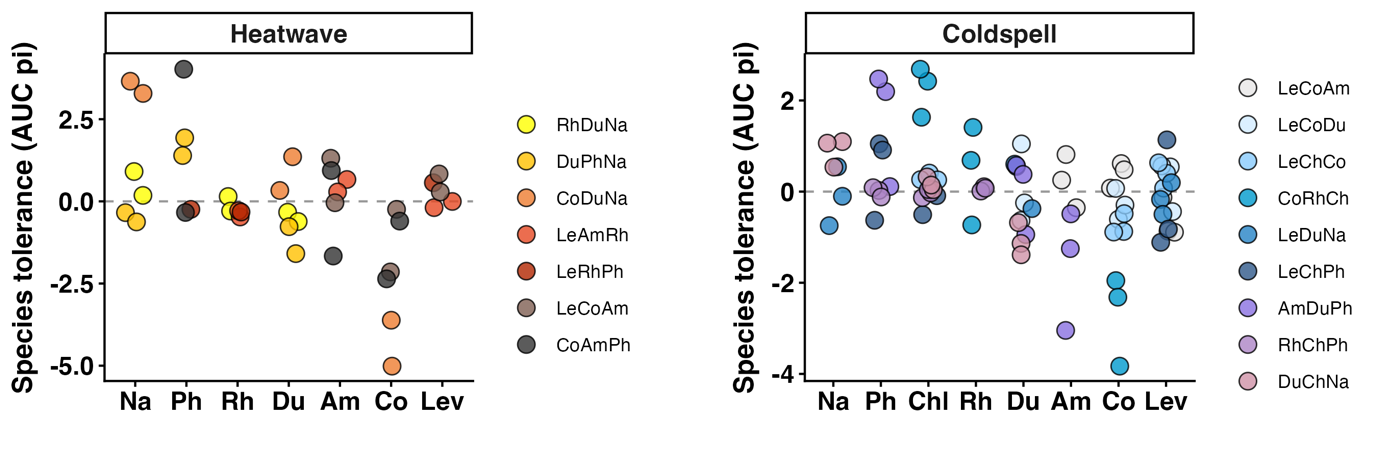

**Figure S7.** Species relative tolerance across communities (differing in imbalance level and composition), including the large species Levanderina. Species on the x-axis are ordered by cell size, from small to large (as shown in Figure 3c species tolerance declines with size for the Heatwave).

**Table S1.** ANOVA outcomes from the linear models testing for the effect of imbalance on stability dimensions across disturbance types. Overall Ecological Vulnerability (OEV) and temporal stability were log10-transformed. For temporal stability, one outlier was removed.

|  |  | Df | Sum Sq | Mean Sq | F-value | P-value |
| --- | --- | --- | --- | --- | --- | --- |
| **Resistance** | Imbalance | 1 | 0.016 | 0.016 | 2.9318 | 0.1 |
|  | Disturbance | 1 | 0.002 | 0.002 | 0.3323 | 0.57 |
|  | Imbalance x Dist | 1 | 0.043 | 0.043 | 7.9799 | **0.008** |
|  | Residuals | 36 | 0.195 | 0.005 |  |  |
| **Recovery** | Imbalance | 1 | 0.052 | 0.052 | 1.8205 | 0.186 |
|  | Disturbance | 1 | 0.0809 | 0.081 | 2.8526 | 0.1 |
|  | Imbalance x Dist | 1 | 0.005 | 0.005 | 0.1882 | 0.667 |
|  | Residuals | 36 | 1.021 | 0.028 |  |  |
| **Resilience** | Imbalance | 1 | 0.004 | 0.004 | 0.162 | 0.69 |
|  | Disturbance | 1 | 0.103 | 0.103 | 4.065 | **0.051** |
|  | Imbalance x Dist | 1 | 0.028 | 0.028 | 1.112 | 0.299 |
|  | Residuals | 36 | 0.913 | 0.025 |  |  |
| **OEV** | Imbalance | 1 | 0.089 | 0.089 | 1.3791 | 0.248 |
|  | Disturbance | 1 | 0.418 | 0.418 | 6.4123 | **0.016** |
|  | Imbalance x Dist | 1 | 0.019 | 0.019 | 0.2857 | 0.596 |
|  | Residuals | 36 | 2.345 | 0.065 |  |  |
| **Temporal Stability** | Imbalance | 1 | 0.006 | 0.006 | 0.1482 | 0.703 |
|  | Disturbance | 1 | 0.024 | 0.024 | 0.6473 | 0.427 |
|  | Imbalance x Dist | 1 | 0.092 | 0.092 | 2.4813 | 0.124 |
|  | Residuals | 35 | 1.332 | 0.037 |  |  |

**Table S2.** ANOVA outcomes from the linear models testing for the effect of species’ traits on species’ sensitivity between disturbance types. The large species Levanderina was excluded.

| Trait |  | Df | Sum Sq | Mean Sq | F-value | P-value |
| --- | --- | --- | --- | --- | --- | --- |
|  | Trait | 1 | 41.14 | 41.14 | 16.14 | **0.0001** |
| ***T*_peak_** | Disturbance | 1 | 0.44 | 0.44 | 0.17 | 0.68 |
|  | Trait x Dist | 1 | 2.68 | 2.68 | 1.05 | 0.31 |
|  | Residuals | 96 | 244.69 | 2.55 |  |  |
| ***µ*max** | Trait | 1 | 81.92 | 81.92 | 40.63 | **<0.0001** |
|  | Disturbance | 1 | 2.79 | 2.79 | 1.38 | 0.24 |
|  | Trait x Dist | 1 | 10.7 | 10.7 | 5.30 | **0.02** |
|  | Residuals | 96 | 193.55 | 2.02 |  |  |
| **Cell size** | Trait | 1 | 63.14 | 63.14 | 26.84 | **<0.0001** |
|  | Disturbance | 1 | 0.001 | 0.001 | 0.0004 | 0.99 |
|  | Trait x Dist | 1 | 0.00 | 0.00 | 0.0002 | 0.99 |
|  | Residuals | 96 | 225.81 | 2.35 |  |  |
| **Size plasticity** | Trait | 1 | 4.72 | 4.72 | 1.61 | 0.21 |
|  | Disturbance | 1 | 0.69 | 0.69 | 0.23 | 0.63 |
|  | Trait x Dist | 1 | 2.23 | 2.23 | 0.76 | 0.39 |
|  | Residuals | 96 | 281.31 | 2.93 |  |  |
| **Dominance** | Trait | 1 | 5.41 | 5.41 | 1.88 | 0.17 |
|  | Disturbance | 1 | 0.01 | 0.01 | 0.004 | 0.95 |
|  | Trait x Dist | 1 | 7.09 | 7.09 | 2.46 | 0.12 |
|  | Residuals | 96 | 276.44 | 2.88 |  |  |

**Appendix 2:**

**# Methods: Species Responses Experiment**

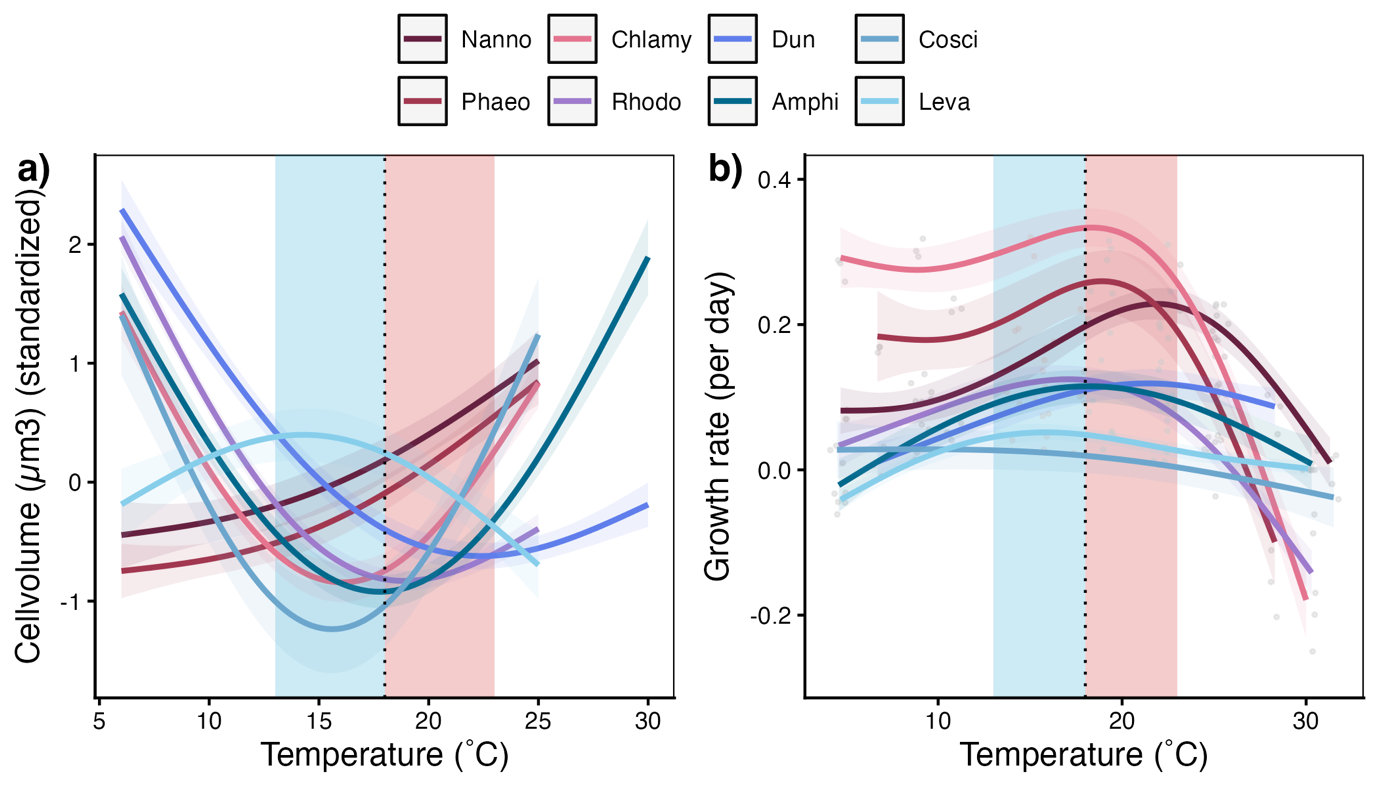

**Figure S8** **a)** Cell size along the temperature gradient during the Species’ Responses experiment to determine cell size plasticity. Blue shaded area presents the temperature range of the size plasticity estimation for the Coldspell (slope from control to Coldspell temperature), and red shaded area the temperature range that was used to calculate the plasticity for the Heatwave (slope from control to Heatwave temperature). **b)** Thermal performance curves measured during the Species’ Responses experiment to determine species’ traits and Imbalance levels for the communities. Coloured shaded areas show the temperature range on what slope estimates are based on for Imbalance calculations (slopes from control to disturbance temperature). Vertical dotted line indicates the control temperature (18˚C) during the disturbance experiment.

**Table S3.** Species traits determined with the Species’ Responses experiment. Cell size was measured before the Species’ responses experiment under culturing conditions. Size plasticity was calculated as the change in cell size (standardized across species) between the control temperature and the disturbance temperature

| **Species** | **Cell size ± SD (µm^3^)** | **Size plasticity** | | | ***T*_peak_ (˚C)** | ***µ*_max_ (day^-1^)** |
| --- | --- | --- | --- | --- | --- | --- |
|  |  | **Heatwave** | | **Coldspell** |  |  |
| *Nannochloropsis* | 17 ± 6 | + 0.13 | | − 0.08 | 22.0 | 0.23 |
| *Phaeodactylum* | 106 ± 95 | + 0.14 | | − 0.09 | 19.0 | 0.26 |
| *Chlamydomonas* | 123 ± 78 | + 0.17 | | + 0.01 | 18.5 | 0.34 |
| *Rhodomonas* | 202 ± 87 | + 0.02 | | + 0.11 | 17.0 | 0.12 |
| *Dunaliella* | 257 ± 80 | − 0.03 | | + 0.18 | 21.5 | 0.12 |
| *Amphidinium* | 713 ± 226 | + 0.1 | | + 0.07 | 18.0 | 0.11 |
| *Coscinodiscus* | 897 ± 228 | + 0.24 | | − 0.02 | 6.5 | 0.03 |
| *Levanderina* | 36 684 ± 14 355 | − 0.15 | + 0.04 | | 16.0 | 0.05 |

**#Methods: Disturbance experiment**:

**Table S4.** Imbalance gradient and species composition in the communities. Crossed-out communities present those that were removed from the experiment due to the incubator that heated up to 28 °C.

| **Coldspell** | |  | **Heatwave** | |
| --- | --- | --- | --- | --- |
| **Imbalance level** | **Community composition** |  | **Imbalance level** | **Community composition** |
| 0.004 | Leva; Cosci;Amphi |  | 0.003 | ~~Leva; Amphi; Nanno~~ |
| 0.006 | Leva; Cosci; Duna |  | 0.004 | Rhodo; Duna; Nanno |
| 0.009 | Leva; Chlamy; Cosci |  | 0.006 | Duna; Phaeo; Nanno |
| 0.011 | Cosci;Rhodo;Chlamy |  | 0.013 | Cosci; Duna; Nanno |
| 0.021 | Leva; Duna; Nanno |  | 0.019 | Leva; Amphi; Rhodo |
| 0.023 | Leva; Chlamy; Phaeo |  | 0.028 | Leva; Rhodo; Phaeo |
| 0.024 | Amphi; Duna; Phaeo |  | 0.028 | Leva; Cosci; Amphi |
| 0.025 | Rhodo; Chlamy; Phaeo |  | 0.037 | Cosci; Amphi; Phaeo |
| 0.032 | Duna; Chlamy; Nanno |  | 0.038 | ~~Leva; Chlamy; Phaeo~~ |

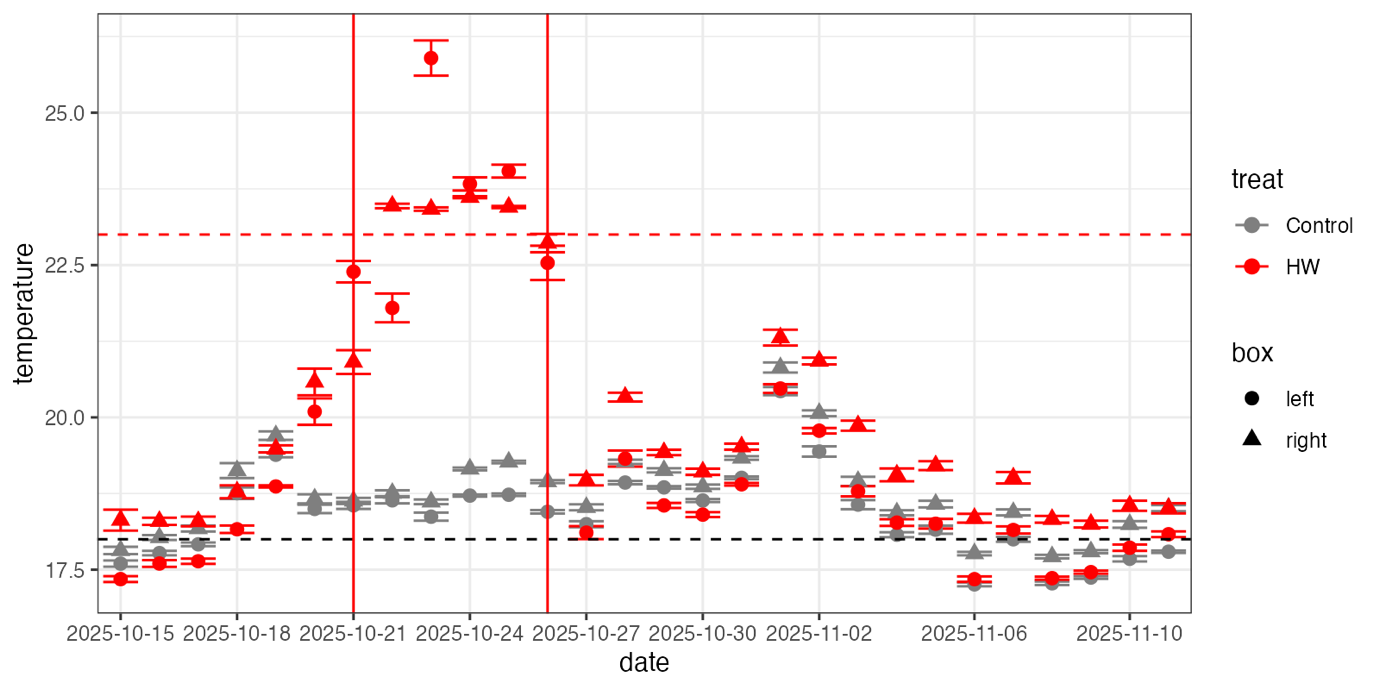

**Figure S9.** Temperature logger data from the Heatwave experiment – averaged for each day and treatment (Disturbance and Control). Dashed horizontal lines present treatment temperature (Control =grey, Heatwave=orange). Solid vertical lines show the window of the Heatwave phase. Shapes show the incubator boxes (for each treatment, two incubator boxes were used).

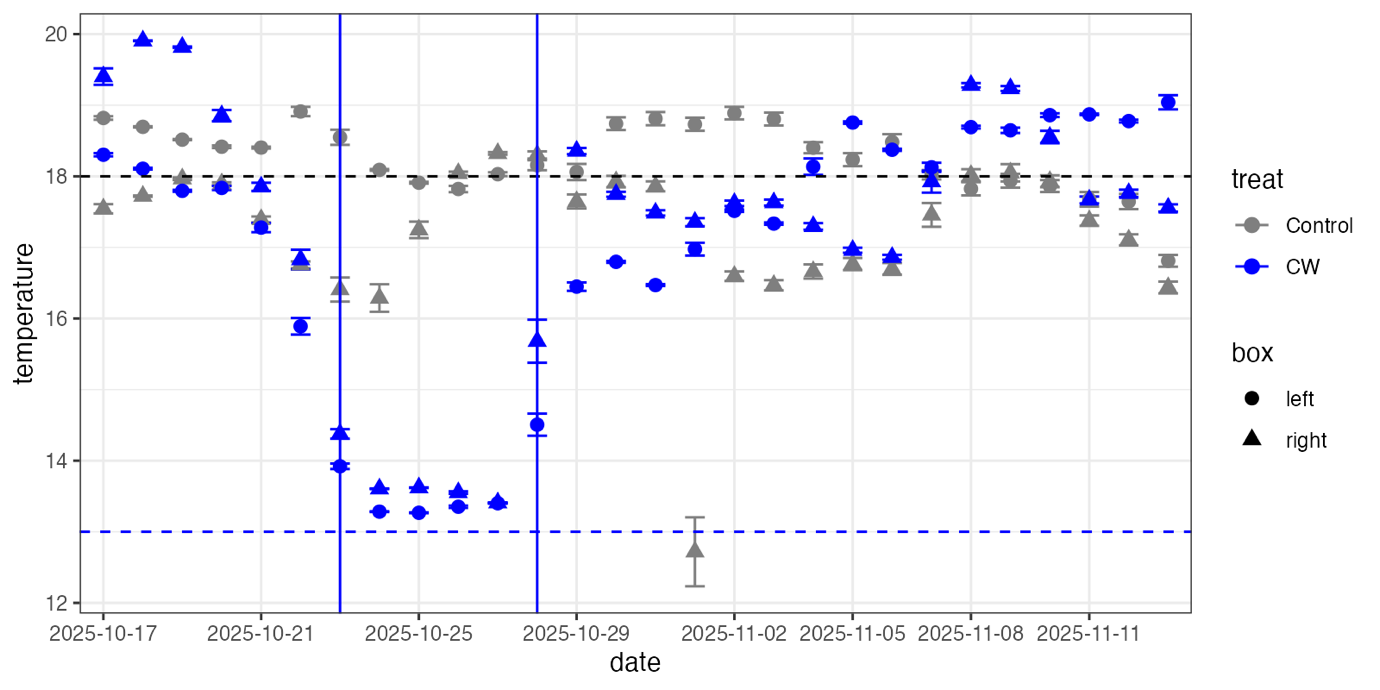

**Figure S10.** Temperature logger data from the Coldspell experiment – averaged for each day and treatment (Disturbance and Control). Dashed horizontal lines present treatment temperature (Control =grey, Coldspell=blue). Solid vertical lines show the window of the Coldspell phase. Shapes show the incubator boxes (for each treatment, two incubator boxes were used).

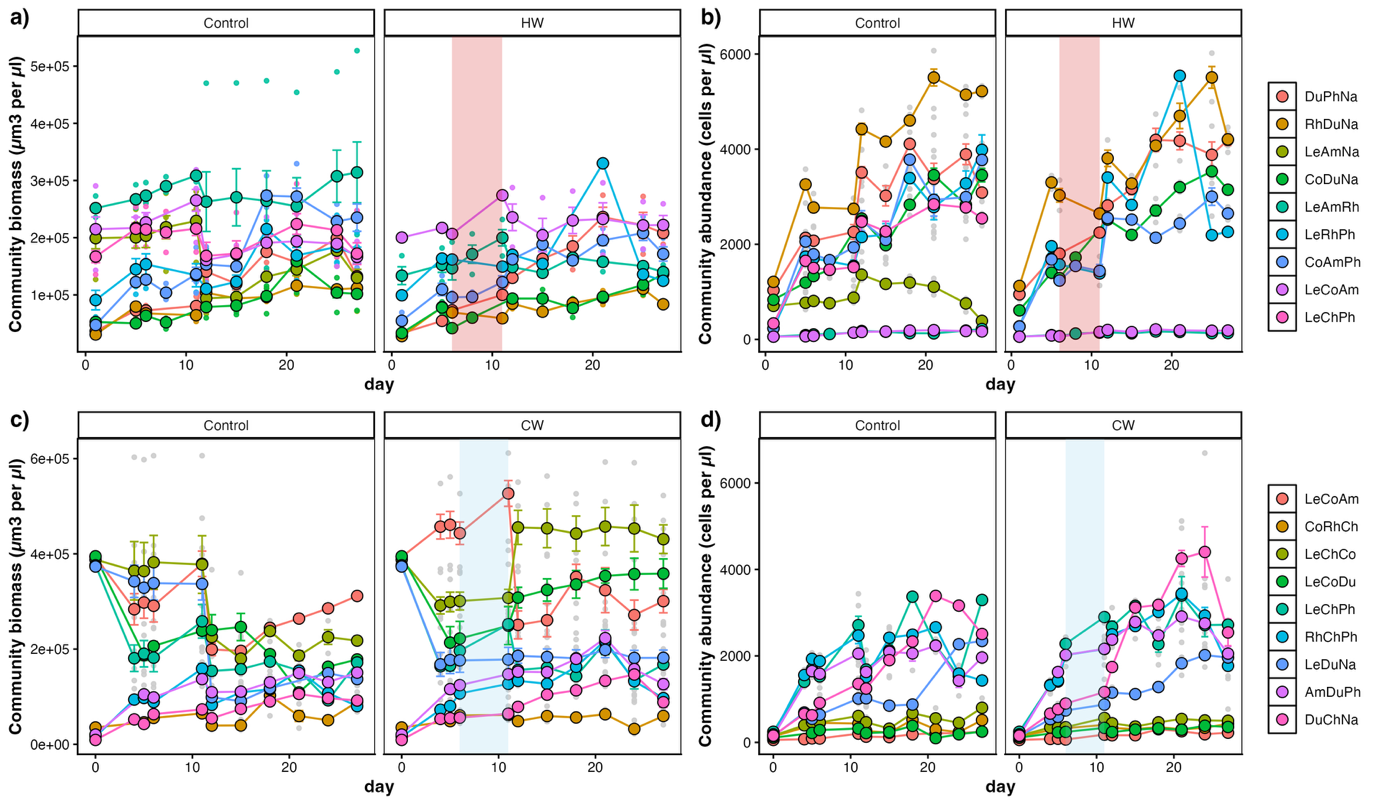

**Figure S11.** Community biomass **(a,c)** and community abundance **(b,d)** over time for each community and treatment (Control and Heatwave, a-b and Coldspell c-d). Replicates are summarized (error bars present standard error). Red and blue shaded areas show the Heatwave and Coldspell phase.

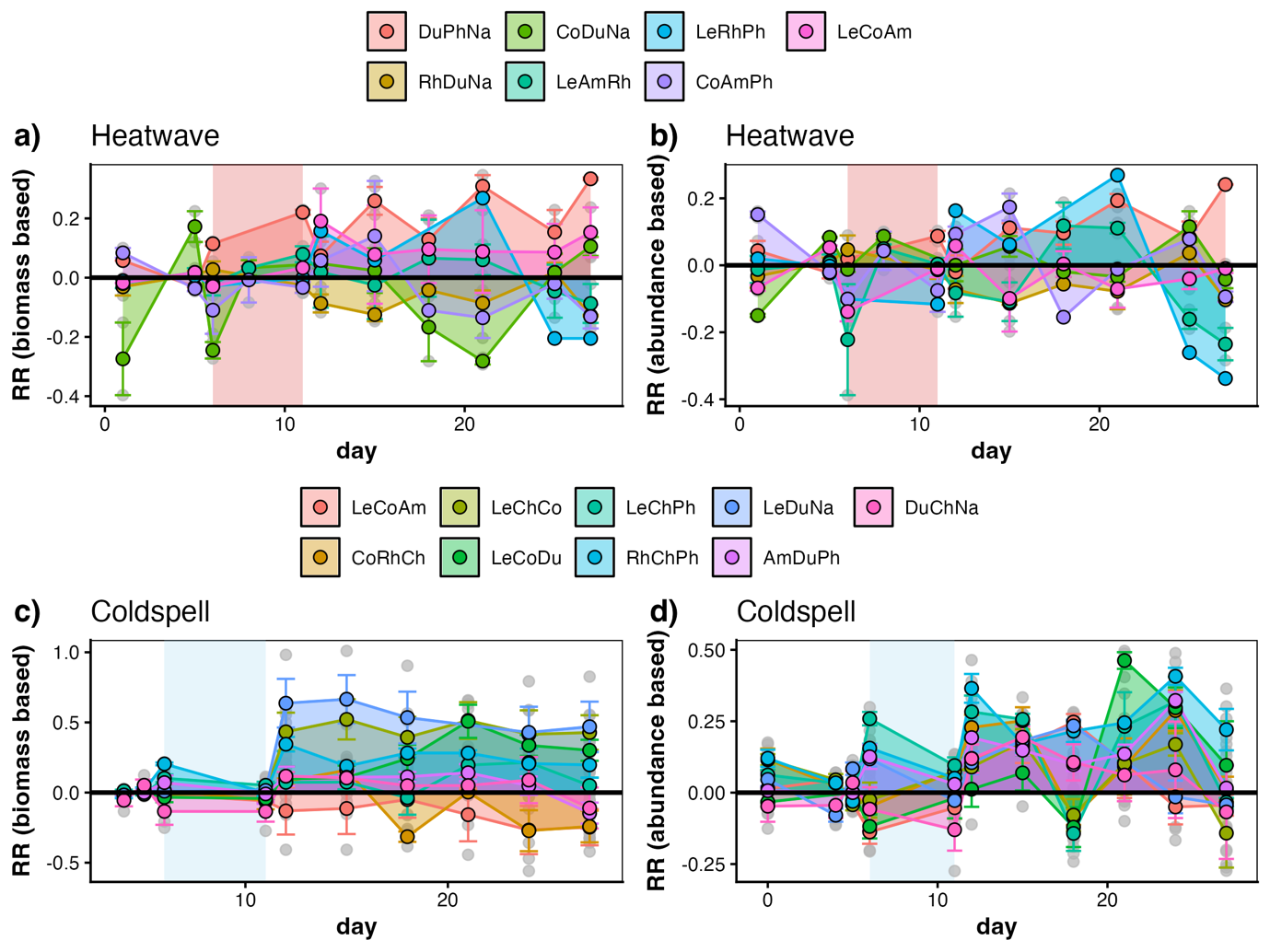

**Figure S12.** Response Ratio (RR) over time of community biomass (a,c) and abundance (b,d) between disturbed communities and control communities, for each disturbance type.
